# Ocean carbon export can be predicted from ocean color-based phytoplankton communities

**DOI:** 10.1101/2024.09.21.613760

**Authors:** Sasha J. Kramer, Erin L. Jones, Margaret L. Estapa, Nicola L. Paul, Tatiana A. Rynearson, Alyson E. Santoro, Sebastian Sudek, Colleen A. Durkin

**Author notes:** Corresponding author: Sasha J. Kramer. **Author Contributions:** SJK, CAD, MLE, TAR, AES designed research. SJK, ELJ, MLE, NLP, AES, SS, CAD performed research. SJK, ELJ, MLE, NLP, SS, CAD analyzed data. SJK and CAD wrote the paper with support from all other authors. The authors declare no competing financial interests.

## Abstract

Carbon flux to the deep sea can be dictated by surface ocean phytoplankton community composition, but translating surface ocean observations into quantitative predictions of carbon export requires additional consideration of the underlying ecosystem drivers. Here, we used genetic tracers of phytoplankton detected in surface seawater and within sinking particles collected in the mesopelagic ocean to identify mechanistic links between surface communities and carbon export in the North Pacific and North Atlantic Oceans. Phytoplankton 18S rRNA sequences were sampled over a one-month period in surface seawater and within bulk-collected and individually-isolated sinking particles using mesopelagic sediment traps (100-500m). Nearly all phytoplankton amplicon sequence variants (ASVs) exported from the surface were packaged in large (>300 µm) particles. Individually, these particles contained only a few distinct phytoplankton ASVs, but collectively, large particles transported about half of the surface taxonomic diversity into the mesopelagic. The relative sequence abundances of the surface community detected within particles were quantitatively related to measured POC fluxes: a linear model based on the relative sequence abundance of just two pigment-based phytoplankton taxa, diatoms and photosynthetic Hacrobia, was predictive of POC flux magnitude. These two taxa were also enriched within the ecologically-distinct particle classes that had the greatest influence on carbon export magnitude. As global, hyperspectral ocean color satellites begin to quantify these taxonomic groups in the surface ocean, the relationship of these taxa to carbon fluxes demonstrated here may help generate more accurate global estimates of export.

## Introduction

Each year, ∼10 Pg of carbon are exported from the surface ocean to the deep sea (1). Best estimates of this component of the global carbon cycle rely on ocean color satellites that collect synoptic observations of surface ocean phytoplankton communities (2). Additional models are required to estimate how much of this primary production is exported from the surface ocean in the form of sinking particles and eventually transported into the deep ocean and sequestered for climate-relevant timescales (3, 4). This process of deep ocean carbon export, known as the biological carbon pump (5), is among the most uncertain components of the global carbon cycle. Identifying mechanisms that link satellite observations of surface ocean phytoplankton with the export of carbon into the deep sea would better constrain the magnitude of the biological carbon pump and enable better predictions of ecosystem responses to changing ocean conditions.

Key processes influencing the biological carbon pump are well described (4), but translating these observations into relationships that are useful for the accurate prediction of carbon export remains challenging. Descriptions of surface ocean phytoplankton community composition have improved through global surveys (e.g., Tara Oceans, Bio-GO-SHIP; (6, 7)) and technologies that increase taxonomic resolution (such as DNA sequencing) and spatial resolution (such as NASA’s Plankton, Aerosol, Cloud, ocean Ecosystem satellite, PACE; (8)). Both the abundance and the taxonomic composition of the surface ocean phytoplankton community can alter the magnitude and efficiency of the biological carbon pump (9, 10). Several studies have identified specific phytoplankton taxa that influence carbon export, such as diatoms and coccolithophores (e.g., (11–15)) or networks of phytoplankton communities associated with elevated carbon flux (16). Other studies have focused on the particles sinking below the surface layer, finding that particles with different ecological origins play distinct roles in packaging surface phytoplankton and transporting carbon into the deep ocean (4, 10).

For example, salp swarms are patchy and episodic, but when salps are present, their fecal pellets efficiently export surface carbon into the deep ocean (e.g., (17)). Collectively, most studies support a key conclusion about the function of the biological carbon pump: not all phytoplankton taxa or particle types are exported with equal efficiency due to variations in cell size and physiology, particle morphology, varying mineral composition, and complex ecosystem interactions such as zooplankton grazing preferences (18, 19). In order to quantitatively link surface phytoplankton communities to the biological carbon pump, we must find a way to mechanistically integrate observations of surface communities with deep sea particle fluxes.

One way to evaluate contributions of different phytoplankton groups to carbon export is to trace their presence in sinking particles in the mesopelagic (19–21). A number of studies have used DNA and RNA sequencing to detect phytoplankton communities within sinking particles captured by sediment traps. Genetic material can be delivered by intact phytoplankton cells or detrital particles, such as zooplankton fecal pellets (e.g., (22–25)). The application of these methods has highlighted the differential contributions of specific phytoplankton taxa to particulate organic carbon (POC) export flux depending on the region and season, as well as the zooplankton grazers that are present to facilitate that export (26–29). Often, high export flux events are comprised of particles containing one or a few dominant taxa (24, 27). Despite these advances, work remains to quantify the predictive power of specific phytoplankton taxa for estimating POC flux into the mesopelagic across the regional and seasonal scales needed to make global estimates. With the advent of the PACE satellite and its Ocean Color Instrument (OCI), it will be possible to observe phytoplankton community composition via pigment-based phytoplankton groups on unprecedented spatial and temporal scales (30, 31). Linking satellite observations to POC flux magnitude and mechanisms requires robust, statistical relationships to be identified among ocean color, phytoplankton community composition, and carbon export flux.

Here, we investigated the distribution of phytoplankton from the surface ocean to the mesopelagic, from whole seawater to sinking particles (19, 32, 33), to answer the following questions: Which phytoplankton taxa are key contributors to carbon export out of the surface ocean? By which mechanisms are these taxa exported? And finally, how can we use this information to build predictive relationships between POC flux magnitude and phytoplankton taxonomy that link to surface ocean color information?

## Results

### DNA-based proxies of surface phytoplankton community composition are comparable to ocean color estimates

To assess whether changes in the surface ocean phytoplankton community detectable by ocean color-based approaches corresponded with signals detectable by DNA sequencing, we compared measured high performance liquid chromatography (HPLC) pigment concentrations, remote sensing reflectance (*R*_*rs*_(λ))-modeled pigment concentrations, and 18S rRNA sequence communities in the surface seawater at two contrasting ocean ecosystems (Figure 1). Variations in the relative abundances of surface phytoplankton groups were generally consistent across observational methods: diatoms were relatively more abundant and photosynthetic Hacrobia were relatively less abundant in the North Atlantic compared to the North Pacific across all methods. By contrast, the relative abundance of all other phytoplankton groups was less consistent, particularly for dinoflagellates. Still, when dinoflagellates were combined with other phytoplankton groups (not including diatoms and Hacrobia) a strong linear relationship existed among observational methods (Figure 1).

**Figure 1.**
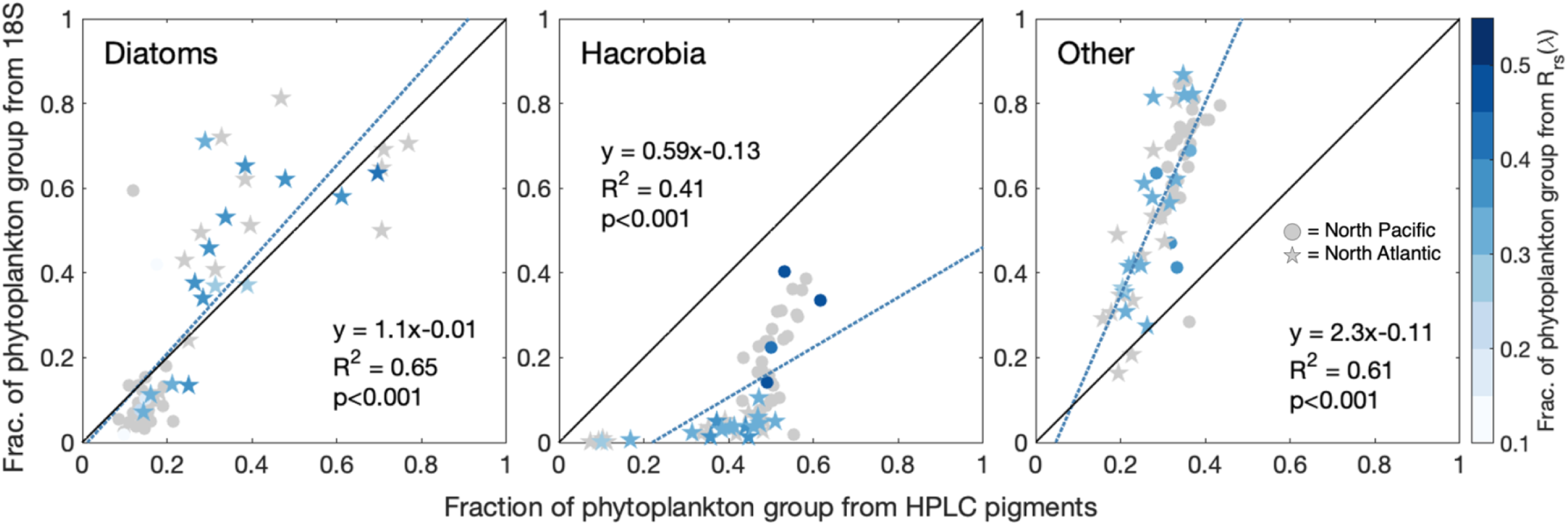
Comparison between surface ocean phytoplankton communities on both EXPORTS field campaigns for major phytoplankton groups examined in this analysis including diatoms, photosynthetic Hacrobia, and other eukaryotic phytoplankton (dinoflagellates, chlorophytes, dictyochophytes, pelagophytes, and other ochrophytes). Circles are North Pacific samples; stars are North Atlantic samples. The relationship between phytoplankton groups as a fraction of all photosynthetic phytoplankton from 18S rRNA gene sequences vs. from measured HPLC pigments is shown, with samples colored by modeled pigments derived from hyperspectral remote sensing reflectances (*R*_*rs*_(λ); [24]) in the cases where those samples were collected at the same places and times; otherwise, samples are grey. Pigment-based phytoplankton groups are represented by: diatoms = Fuco, photosynthetic Hacrobia = HexFuco + Allo, other = Perid + MVchlb + ButFuco + 20% of Chlc.

### Composition and diversity of phytoplankton varies between basins across surface seawater and sinking particles

Phytoplankton communities differed by ocean basin, depth, and collection method based on 18S rRNA gene relative abundances. All North Pacific samples differed significantly from all North Atlantic samples (Figure 2A; PERMANOVA of Aitchison distances computed from PCA; adjusted p = 0.006). North Atlantic samples were positively associated with diatoms, especially the genera *Chaetoceros*, *Thalassiosira*, and *Pseudo-nitzschia*. North Pacific samples were positively associated with dinoflagellates (*Gymnodinium*, *Gyrodinium*, *Prorocentrum*), Hacrobia (*Phaeocystis* [prymnesiophyte], *Chrysochromulina* [prymnesiophyte], *Plagioselmis* [cryptophyte]), and chlorophytes (*Chloroparvula*, *Bathycoccus*; Figure 2B). The largest variability among all samples was between surface seawater communities and all other sample types, which separated on the first principal component (26.5% of variability; Figure 2A; PERMANOVA of Aitchison distances; NP adjusted p = 0.003; NA adjusted p = 0.003). Communities found in individual particles also differed from those found in bulk particle samples (PERMANOVA of Aitchison distances; NP adjusted p = 0.0028; NA adjusted p = 0.0021).

**Figure 2.**
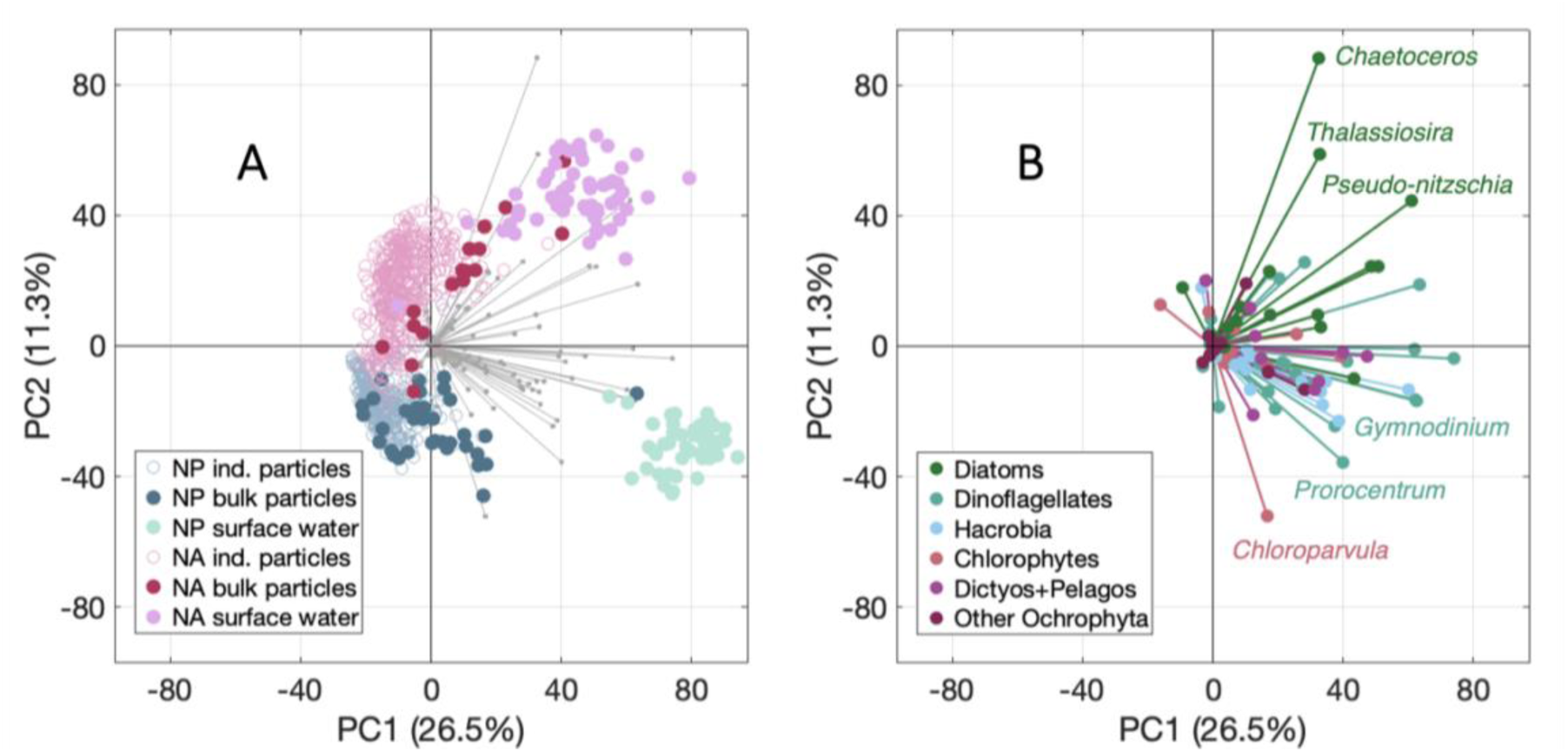
Principal components analysis plot showing the first two components for all surface, bulk trap, and particle samples in both basins ((A) North Pacific in blues and North Atlantic in pinks) and genus-level ASV bins driving this variability (grey vectors in (A) and colored vectors in (B)). A subset of the genera with the largest loadings on each axis are labeled in (B).

Despite their differing compositions, samples of the same type shared similar alpha diversity of phytoplankton (amplicon sequence variant [ASV] richness) between the two ocean basins (Figure 3). The fraction of ASVs shared among all three sample types (surface seawater, bulk particles, and individual particles) was small (North Pacific = 105 ASVs, 8% of total; North Atlantic = 151 ASVs; 12% of total). The surface seawater communities in the North Pacific were more diverse (526 ASVs) than in the North Atlantic (410 ASVs). A greater proportion of that diversity was only found in the surface seawater (274 ASVs; 21% of total) compared to the North Atlantic (150 ASVs; 12% of total). In contrast to the average surface seawater sample, the average diversity within individual particles was low (Figure 3, inset boxplots), with a slightly higher per-particle diversity detected in the North Atlantic (median 14 ASVs per particle) than the North Pacific (median 8 ASVs per particle). However, when considered all together, the particle pool contained the greatest fraction of total diversity, most of which was not shared with other sample types. In the North Pacific, individual particles comprised 584 ASVs (or 44% of the total) that were not found in the other pools. In the North Atlantic, individual particles comprised 595 ASVs (or 49% of the total) that were not found in other pools. In each basin, the mean ASV diversity in each sample type differed significantly from the other two sample types (one-way ANOVA with Tukey-Kramer posthoc; NP p<0.001; NA p<0.001). Each individual particle transported a small and highly variable subset of the total phytoplankton diversity found in the bulk sinking particles and the surface seawater. The bulk sinking particle communities had the lowest number of unique ASVs (159 or 12% in the North Pacific; 94 or 7% in the North Atlantic).

**Figure 3.**
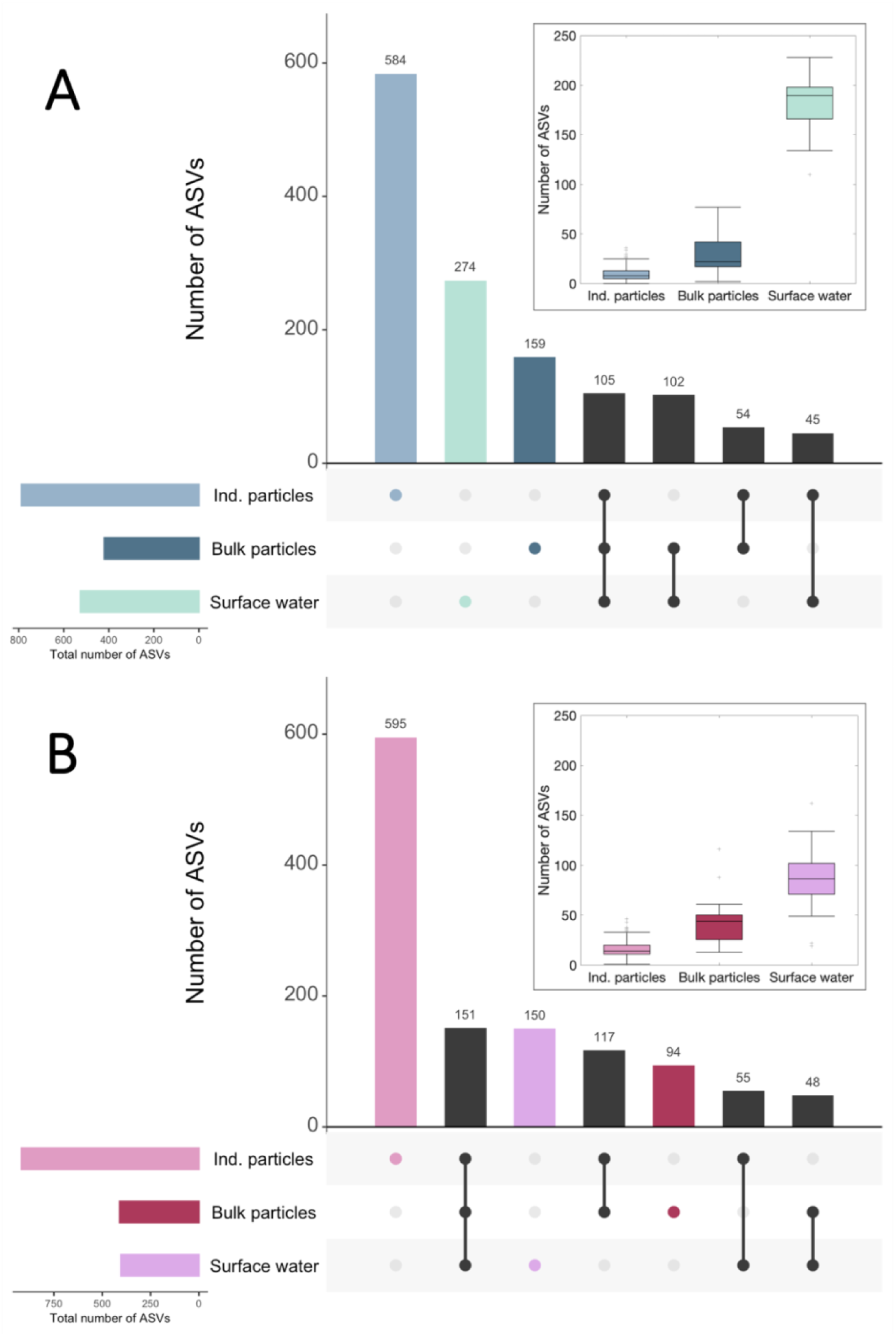
Number of phytoplankton ASVs shared among three sample types in (A) the North Pacific and (B) the North Atlantic. Bars in the upper panel show the number of unique or shared ASVs in each sample category or grouping (indicated by the dots in the lower panel). Bars on the left panel show the total number of ASVs in that sample category. Inset boxplots show the average number of ASVs per sample across all samples in each category (center line) and the range of values to the 25^th^ and 75^th^ percentiles, respectively (edges of the box). Whiskers cover +/- 2.7 times the standard deviation of the data and outliers from this range are shown as grey crosses.

### Most exported surface phytoplankton taxa are packaged in large sinking particles

In the North Pacific, a little less than half of the surface ASV diversity was detected in sinking particles on average, while in the North Atlantic, more than half of the average surface phytoplankton ASV diversity was also found in sinking particles on average (summed bulk and individual particles; Figure 4). Most exported phytoplankton ASVs from the surface were detected in large (>300 μm), individual sinking particles rather than only in bulk-collected particle samples. While there were major differences in sinking particle abundances and size distributions in the North Pacific (32, 34) vs. the North Atlantic (35), these large particles remained the primary export mechanism for surface phytoplankton ASVs in both basins (Figure 4). A similar link between large particles and surface phytoplankton was identified when the comparison was reversed (all phytoplankton ASVs in sinking particles vs. those shared in the surface seawater, Figure S2). Finally, the majority of the ASVs found in bulk sinking particles were also shared with individual particles (Figure S3), further demonstrating the dominance of larger particles for packaging phytoplankton ASVs.

**Figure 4.**
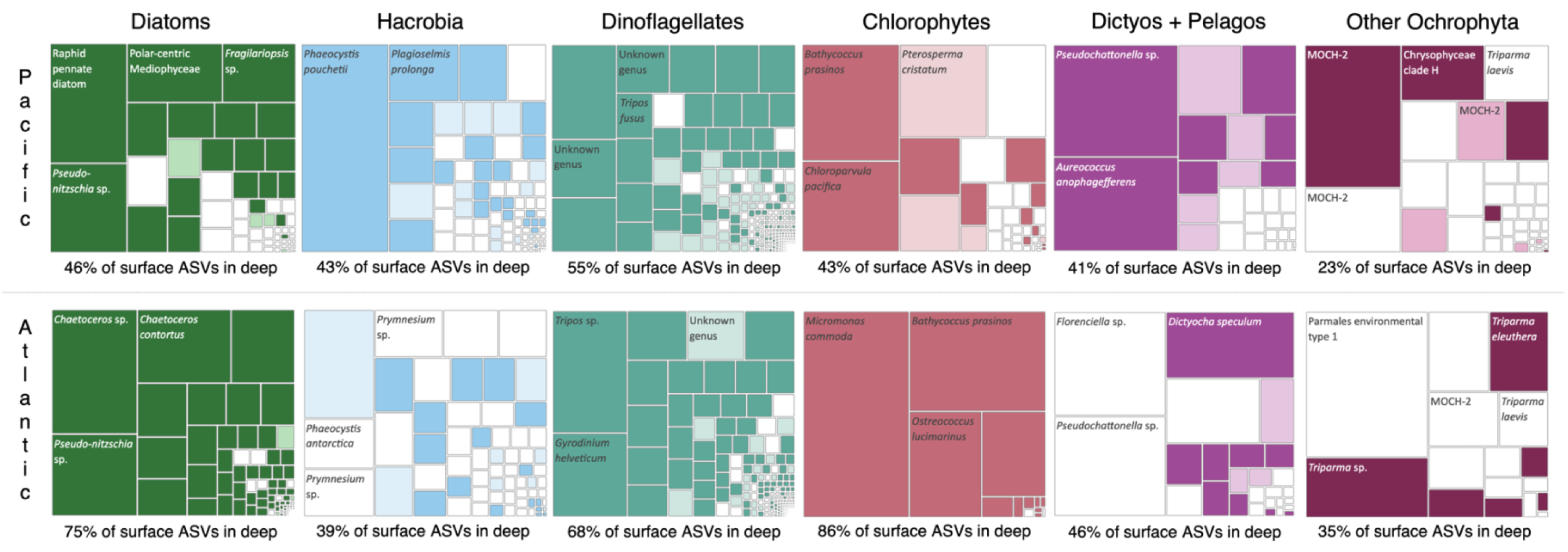
Major pigment-based phytoplankton group abundance in surface seawater in the North Pacific (top row) and North Atlantic (bottom row). Each small box within a larger box represents an individual ASV within that group. If the box is colored in, that ASV was found in a sediment trap sample (darker color = in bulk particles and individual particles, lighter color = only in bulk particles, not individual particles). Boxes that are not colored in represent ASVs that were not found in sinking particles.

### Diatom and photosynthetic Hacrobia enrichment in particles of differing ecological origin

Given that most exported phytoplankton ASVs were found in large particles (Figure 4), we next investigated if the phytoplankton composition within individual particle classes was distinct. The contribution of each particle type to POC flux was established using quantitative imaging (33), to compare the magnitude of flux in each particle class to the phytoplankton community within that particle class from 18S rRNA gene sequences (Figure 5). In the North Pacific, the composition of all individual particles was statistically similar across particle types, with the exception of salp fecal pellets (Figure S4; PERMANOVA of Aitchison distances; adjusted p = 0.0049). When ASV relative abundances were summed together by type, salp fecal pellets contained higher relative diatom and Hacrobia sequence reads on average than other particle types (Figure 5), despite the high variability within individual particle samples (Figure S4). In the North Atlantic, three ecologically distinct particle classes contained different sequence compositions: aggregates and dense detritus, large loose pellets and long fecal pellets, and short pellets (PERMANOVA of Aitchison distances; adjusted p = 0.0021). The relative diatom sequence abundance was 15-25% higher on average in the first two particle categories compared to other North Atlantic particle types (Figure 5).

**Figure 5.**
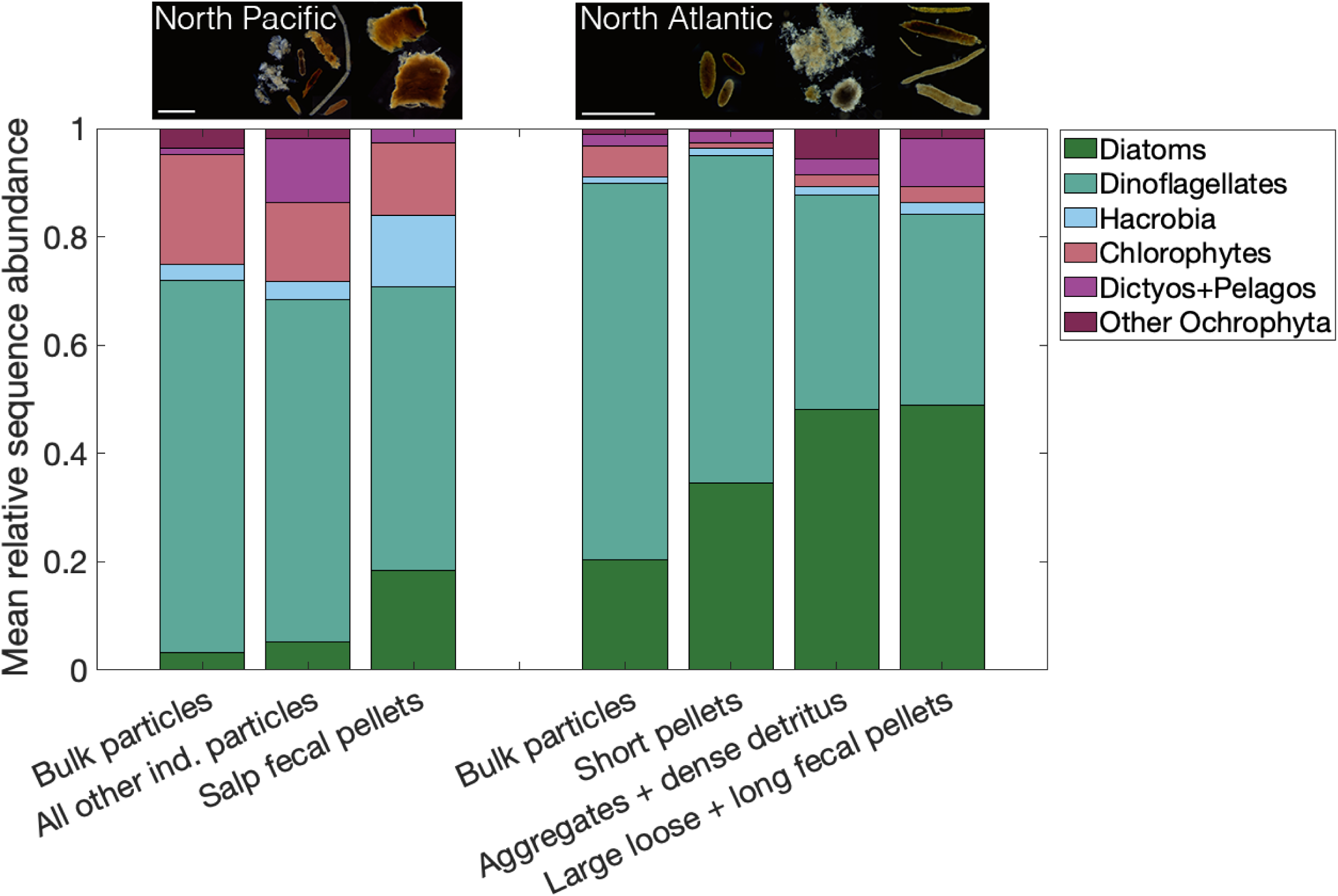
Mean relative sequence abundance of major pigment-based phytoplankton taxa within bulk particles and summed individual particles from statistically different particle types in both basins. Dark-field microscopy images of major particle types are shown above the corresponding bar for each basin (white scale bar = 1 mm).

### Major phytoplankton taxa enriched in large sinking particles are predictive of POC flux magnitude in bulk traps

The enrichment of specific taxa in the large sinking individual particles was directly related to variability in POC flux through PCA of bulk particle sample DNA communities (Figure 6A). The first component (37.0% of variability) was most strongly negatively correlated with POC flux (R^2^ = 0.83) and the relative sequence abundances of diatoms (R^2^ = 0.38) and Hacrobia (R^2^ = 0.48), and positively correlated with the relative sequence abundance of dinoflagellates (R^2^ = 0.88). The second PCA component (21.5% of variability) was strongly positively correlated with the relative sequence abundances of Dictyochophytes + Pelagophytes (R^2^ = 0.71) and other Ochrophyta (R^2^ = 0.76). Twenty six of 28 North Pacific samples were associated with a positive Component 1 (higher relative sequence abundances of dinoflagellates) and 14 of 15 North Atlantic samples had negative loadings on Component 1 (higher relative sequence abundances of diatoms and Hacrobia; higher POC flux). This relationship was reflected on the individual particle level (Figure S5).

**Figure 6.**
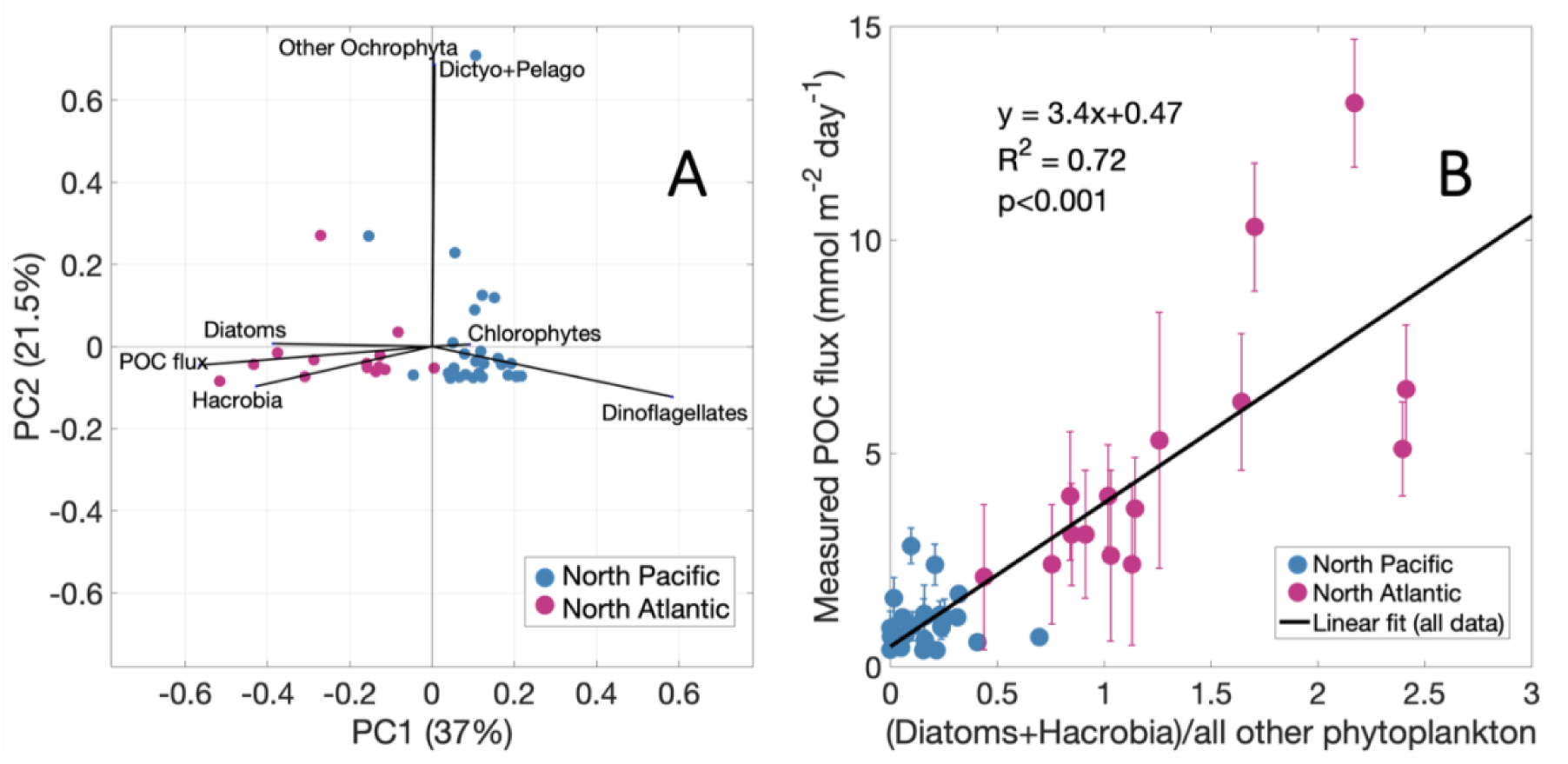
(A) Principal components analysis of pigment-based phytoplankton groups obtained from ASV data in bulk sediment traps. The orientation of each sample (North Pacific in blue, North Atlantic in pink) and each variable (black vectors) are shown for the first 2 components. (B) Relationship between POC flux and the ratio of diatoms + Hacrobia to all other photosynthetic eukaryotic phytoplankton detected within bulk sinking particles. Samples are colored by ocean basin (North Pacific in blue, North Atlantic in pink). Error bars represent the uncertainty associated with each POC measurement. The linear fit is shown in black.

We tested a simple POC flux model based on the ratio of diatom and Hacrobia sequences to all other phytoplankton sequences detected in bulk sinking particles. We calculated a linear relationship (R^2^ = 0.72, p<<0.001) using the relative abundance of only ASVs that were shared between sinking particles and surface seawater (Figure 6B; Figure S2). No significant relationship was found when performing this analysis on all phytoplankton taxa detected in particles but not in surface ocean samples. In the North Atlantic, sinking particles exported more total POC and contained greater relative sequence abundances of diatoms and Hacrobia relative to all other phytoplankton, while in the North Pacific, the POC flux was lower, with corresponding higher relative abundances of other phytoplankton in sinking particles relative to diatoms and Hacrobia (Figure 6B). There was no independent linear relationship found between these variables in only the North Pacific samples (y = 0.17x + 0.97; R^2^ = 0.002, p = 0.83) and a weaker relationship found in only the North Atlantic (y = 3.4x + 0.24; R^2^ = 0.48, p = 0.003).

Although the linear relationship was not as strong within an individual basin, the linear model still adequately represented the spatial and temporal variability in POC flux detected at each site over the course of one month (Figure 7). In the North Pacific, dinoflagellate reads dominated the phytoplankton community within sinking particles, comprising 62-98% of sequence reads in Deployment 1 and 2 (Figure 7A and 5C). In Deployment 3, however, the relative abundance of diatom reads increased at 95m and 195m (Figure 7E), corresponding to higher POC flux during that time (Figure 7E-F).

**Figure 7.**
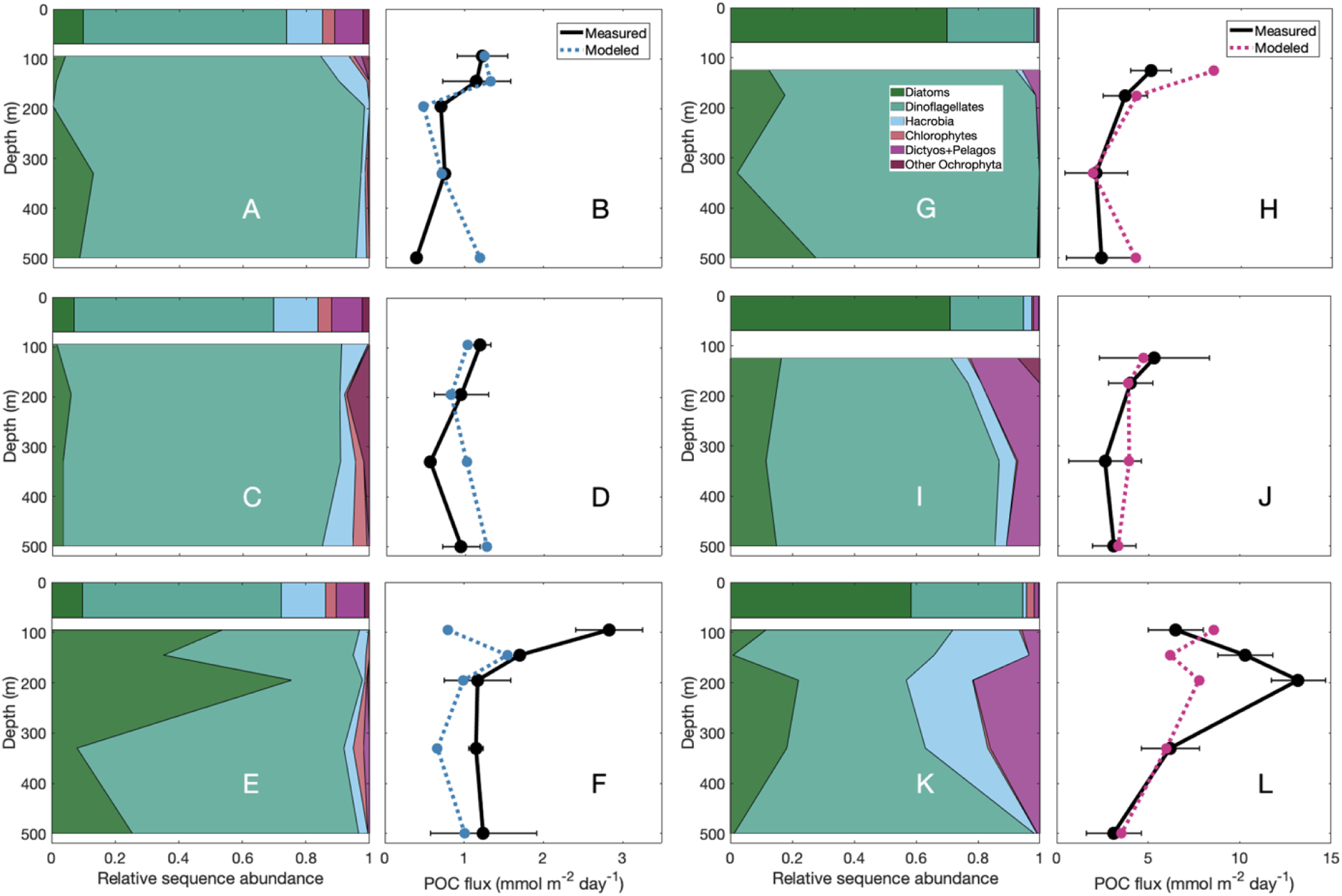
Phytoplankton community composition determined from 18S relative sequence abundance, measured POC flux, and modeled POC flux from the (diatoms + Hacrobia) / all other phytoplankton ratio (by applying the equation in Figure 6B), all from bulk particle samples, over space and time. Samples are shown in the North Pacific deployment 1 (A-B), deployment 2 (C-D), and deployment 3 (E-F) and in the North Atlantic deployment 1 (G-H), deployment 2 (I-J), and deployment 3 (K-L). All samples shown here were collected from surface-tethered traps (STTs) to maximize match-ups between POC and 18S rRNA samples across depths and deployments.

POC flux was typically low in the North Pacific (0.5-1.5 mmol m^-2^ day^-1^), with the exception of the 95m trap in Deployment 3 (2.8 mmol m^-2^ day^-1^). To compare measured POC flux to that predicted using DNA sequence data, we used the line of best fit to model POC flux from the ratio of phytoplankton relative sequence abundances (Figure 6B; y = 3.4x+0.47, where x = [diatoms + Hacrobia]/all other phytoplankton). The modeled values varied over the 3 deployments in the North Pacific and broadly mirrored the shape and magnitude of the measured POC flux depth profile down to 500 m (Figure 7B, 7D, 7F).

In the North Atlantic, the surface seawater contained relatively higher diatom sequence reads than the North Pacific, comprising 70% of the reads in Deployment 1 (Figure 7G), 71% in Deployment 2 (Figure 7I), and 59% in Deployment 3 (Figure 7K). There were also higher relative sequence abundances of Hacrobia, dictyochophytes, and pelagophytes, particularly in Deployment 2 and 3 (Figure 7I, 7K). These phytoplankton communities corresponded to higher POC flux values that increased from Deployments 1 and 2 (2.1-5.3 mmol m^-2^ day^-1^; Figures 7H, 7J) to Deployment 3 (6.2-13.2 mmol m^-2^ day^-1^; Figure 7L). The modeled values again demonstrated the relationship between the phytoplankton community (in this case, higher relative sequence abundances of diatoms and Hacrobia) and higher POC flux (Figures 7H, 7J, 7L).

## Discussion

We identified a strong correspondence between phytoplankton community composition and POC flux magnitude as measured in bulk sediment traps using a predictive linear model that incorporated 18S rRNA gene sequences within sinking particles. Notably, the two key taxa, diatoms and photosynthetic Hacrobia, are detectable using ocean color remote sensing by modeling phytoplankton pigment concentrations. The variation in relative phytoplankton pigment contributions detected among remotely-sensed phytoplankton groups was comparable to the variations in relative abundances observed using 18S rRNA sequencing in surface seawater.

Although the resulting predictive relationship between 18S-based phytoplankton community composition in sinking particles and POC flux was identified using basin-scale variability, the relationship also reasonably predicted POC fluxes within both basins with depth and over time. These results support previous studies that identified links between diatoms and some Hacrobia taxa (specifically prymnesiophytes such as coccolithophores) and POC flux (e.g., (13, 16)). By probing the contents of hundreds of individually-isolated particles, we further identified some of the mechanisms driving these correlations: these key taxa were preferentially packaged in large particles and in particles with ecological origins that contributed the most to carbon flux variability in each ecosystem (e.g., salp fecal pellets, aggregates; (17, 35)).

About 50% of surface ocean phytoplankton ASVs were also found in sinking particles, and reads from these shared taxa tended to be more relatively abundant in the surface. The shared taxa had a quantitative relationship with POC flux when packaged in sinking particles, suggesting a specific link between POC flux and surface ocean conditions. While sinking particles contained ASVs not detected in the surface ocean, isolating the taxa that were shared between the surface seawater and sinking particles was the only approach that effectively predicted POC flux at those trap depths. These genetic tracers of the surface phytoplankton community predicted 72% of the bulk POC flux. The distribution of phytoplankton ASVs across particle sizes and types illuminated the mechanisms driving POC flux, particularly phytoplankton packaging.

While only a few phytoplankton ASVs were packaged into an individual particle, together these large (>300μm) particles contained most of the exported surface phytoplankton diversity. In general, the surface taxa with the most abundant reads were recovered in sinking particles. Specific types of large sinking particles were important contributors to POC flux during both EXPORTS field campaigns (e.g., salp fecal pellets in the North Pacific (17), aggregates in the North Atlantic (35)). Importantly, key phytoplankton taxa that positively correlated with POC flux (diatoms and photosynthetic Hacrobia) had relatively more abundant reads in these particle types.

While each method of describing surface ocean phytoplankton community composition has its strengths and weaknesses (e.g., see (36) for an example using the EXPORTS North Pacific dataset), these methods are strongest when they can be combined to extend the utility of one method beyond its limitations. For instance, in this study, gene sequencing data was needed to achieve higher taxonomic resolution and compare between surface seawater samples and sinking particle samples, while phytoplankton pigment-based taxonomy was required to translate the information in sequencing data to something remotely-sensible by ocean color satellites. The relationships between phytoplankton community composition methods found here persist despite notable challenges in comparing between these approaches. Not all dinoflagellates identified by 18S rRNA are photosynthetic, and not all photosynthetic dinoflagellates contain peridinin (the dinoflagellate biomarker pigment; (37)). Further, 18S rRNA sequencing may overestimate the relative abundances of dinoflagellates due to high copy numbers of the 18S rRNA gene (38). Importantly, all three methods are needed to characterize the phytoplankton community: pigments modeled from hyperspectral *R*_*rs*_(λ) link ocean color to surface ocean phytoplankton, in situ HPLC pigments ground-truth the satellite measurements and can be compared to the surface 18S rRNA samples, and 18S rRNA enables us to trace surface phytoplankton in sinking particles. Together, these methods translate phytoplankton community composition from ocean color to the surface seawater and into deep sinking particles. The ecosystems investigated here encompass a high dynamic range in phytoplankton community composition and POC flux conditions, but the relationships we identified among pigments, DNA, and POC flux will need to be tested on broader global scales and using phytoplankton information at varying taxonomic resolution.

In this analysis, we demonstrated agreement between disparate methods. Ocean color satellites view only the surface layer, and the magnitude of sinking POC or the mechanisms by which it is exported are largely invisible from space. However, by developing a predictive ratio of pigment-based phytoplankton taxa shared between the surface and deep ocean to model POC flux in the mesopelagic, we show that there are quantitative links between phytoplankton community composition, POC export flux magnitude, and mechanisms for export. This work starts to identify which information from the surface ocean is needed from satellites to predict POC flux dynamics in the mesopelagic. We suggest that diatoms and Hacrobia, which can be detected from surface ocean phytoplankton pigments and thus from PACE-resolution data, are particularly important. Since PACE launch in February 2024, the available information about surface ocean color and phytoplankton communities around the globe is rapidly increasing. With this improved resolution to describe the surface ocean phytoplankton community, we will be able to extend the predictive model shown here to the surface ocean and test its performance across ecosystems.

## Materials and Methods

### Overview of contrasting ecosystems

Sampling was conducted as part of the EXport Processes in the Ocean from RemoTe Sensing (EXPORTS) field campaigns aboard R/V *Roger Revelle* in the eastern North Pacific Ocean (Ocean Station Papa, August-September 2018; (39)) and aboard RRS *James Cook* in the eastern North Atlantic Ocean (Porcupine Abyssal Plain, May 2021; (40)). Sampling included sediment trap deployments and whole seawater sample collection. POC flux magnitude was higher in the North Atlantic, driven by pulses of fresh phytodetritus and intense storm activity (41–43). In the North Pacific, a highly recycled ecosystem was sampled, with lower POC flux magnitude that was at times enhanced by salp grazing activity (17, 34, 43, 44).

### Sediment trap sampling

On each cruise, sediment traps were deployed three times, for a duration between 2 to 6 days. Details of sediment trap deployments are found in (34) and (45). DNA and POC samples described here were collected from both Surface-Tethered Traps (STTs; (46)) and Neutrally Buoyant Sediment Traps (NBSTs; (47)). The STT array collected sinking particles at five depths while the NBSTs were deployed at a subset of those five depths. The shallowest traps were deployed below the euphotic zone, defined by the 1% surface light level. Traps were deployed in three successive collection periods over one month. Additional details about sediment trap preparation and deployment are found in the Supporting Information.

### Bulk sediment trap samples

Bulk particle collections (North Pacific n = 35; North Atlantic n = 17) were sampled for nucleic acids following (48). Particles from one collection tube were filtered with vacuum filtration onto a 0.2 μm Supor filter, placed in cryovials, flash frozen in liquid nitrogen, and stored at −80°C. Total DNA was extracted using the PowerViral DNA/RNA extraction kit (Qiagen, Hilden, Germany) following manufacturer’s instructions. These samples are described as “bulk particles.”

### Individual particles

Imaging of particles collected in polyacrylamide gels is described in (32) and the collection of nucleic acids from individual sinking particles is described in (19, 48). Polyacrylamide gels were removed from the trap tubes upon recovery and quantitatively imaged at multiple magnifications using a stereomicroscope (Olympus SZX16 with Lumenera Infinity 2 camera attachment). Imaged particles were quantified, sized, and classified using Python-based image processing scripts and their contributions to POC flux were modeled following (32, 33). After imaging, particles larger than ∼300 μm (North Pacific n = 320, North Atlantic n = 450) were isolated from the gel using a 1000 μL micropipette and transferred into a cryovial, where they were flash frozen in liquid nitrogen and stored at −80°C until analysis. Four major particle categories were separated via imaging: 1) salp fecal pellets, 2) aggregates and dense detritus, 3) long fecal pellets and large loose pellets, and 4) short pellets (Figure S1). Individual particle DNA was extracted in a 5-10% solution of Chelex 100 resin (Bio-Rad, Hercules, CA, USA) by iteratively incubating (at 95°C) and vortexing the sample until solid debris was concentrated at the bottom of the sample tube via centrifugation. The overlying liquid containing nucleic acids was removed (19). The DNA was further purified and concentrated using the DNA Clean and Concentrator kit (Zymo Research, Orange, CA, USA).

### Euphotic zone seawater sampling

Whole seawater samples were collected in Niskin bottles from three depths (surface, 0.1% light, and 0.01% light) at noon during days spanning the duration of the trap deployments in both the North Pacific and North Atlantic. Samples were filtered onto 0.2 µm pore size, 47 mm polyethersulfone filters (Express Plus; Millipore), flash frozen in liquid nitrogen, and stored at −80 °C until processing. Total DNA was extracted from plankton biomass samples (North Pacific n = 54, North Atlantic n = 54) using the Quick DNA/RNA Miniprep Plus Kit (Cat. No. D7003; ZymoResearch) following manufacturer’s instructions for tissue extraction with the following modifications: 150 μl 0.4mm Zirconium beads, 60µl PK Digestion Buffer, and 30 µl Proteinase K were added prior to homogenization (30 secs), incubation (2 hours at 55°C), vortexing, and centrifugation to pellet debris (2 mins maximum speed).

### Analysis of DNA sequences

Details of 18S rRNA gene amplification and sequence analysis are found in the Supporting Information. Processing of 18S rRNA gene sequence reads for all sample types (seawater, bulk traps, and individual particles) was conducted as described in (32) using QIIME2 (ref 49) and the DADA2 (ref 50) workflow. Taxonomy was assigned to ASVs using a naïve Bayesian classifier algorithm (51) with a minimum bootstrap confidence of 80% for the Protist Ribosomal Reference ((52); PR2 v 4.14.0, downloaded September 2022). ASVs were assigned as either photosynthetic or heterotrophic taxa in accordance with prior trophic classification by (19) and with supplemental photosynthetic taxa unique to this dataset. Our analysis focused on photosynthetic phytoplankton ASVs. Dinoflagellate trophic strategies are variable and often fluid, so we included all dinoflagellates in our analysis of phytoplankton while acknowledging that some taxa may be heterotrophic. Known parasitic dinoflagellate taxa (e.g., *Syndiniales* spp.) were removed. Based on ASV accumulation curves, samples reached saturation at 5,000 reads per sample. Thus, we removed 9 samples with <5,000 reads from further analysis. The code for these analysis steps, including the list of photosynthetic taxa, is available on GitHub (see Data Availability Statement).

Phytoplankton ASV presence and relative abundance was compared across sample types. Taxa identified in both the individually picked particles and corresponding bulk sediment trap samples from the same trap platform were considered to be packaged within “large” (>300 μm) particles. Taxa found in the bulk sediment trap samples but not in the individual particles isolated from the same trap platform were assumed to be packaged in particles smaller than could be isolated for successful DNA amplification (<300 μm).

Relative phytoplankton read counts were center log-ratio transformed and sample composition was compared using a compositional principal components analysis (PCA) in Matlab v. R2022a (53). The singular value decomposition (SVD) approach was used to construct the PCA (“svd” function in Matlab) and photosynthetic phytoplankton sequences were grouped to the genus-level for this analysis (N = 132). Differences among sample groups were assessed from Aitchison distances (54) using PERMANOVA with Bonferroni p-value adjustment for multiple comparisons. PCA was also used to examine the co-variability in phytoplankton groups and POC flux. For this analysis, SVD was not used to calculate the PCA (“pca” function in Matlab). Variables were mean-centered and normalized by their standard deviation before PCA was performed.

### Phytoplankton community composition from other methods

For all further analyses, phytoplankton taxa from 18S rRNA were divided into six pigment-based taxonomic groups (following (55)). The phytoplankton groups and their assumed corresponding biomarker pigments are as follows: diatoms (fucoxanthin), dinoflagellates (peridinin), photosynthetic Hacrobia (prymnesiophytes [19’hexanoyloxyfucoxanthin] and cryptophytes [alloxanthin]), Chlorophytes (monovinyl chlorophyll b), Dictyochophytes and Pelagophytes (19’butanoyloxyfucoxanthin), and other Ochrophyta (20% of total chlorophyll c, assuming an equal distribution across all red algal classes shown here). Phytoplankton pigment concentrations measured by high performance liquid chromatography (HPLC) from each EXPORTS field campaign were compared to modeled pigment concentrations from hyperspectral remote sensing reflectance (*R*_*rs*_(λ); sr^-1^) generated using the Spectral Derivatives Pigment (SDP) model (30). Details for HPLC pigment sample collection and hyperspectral *R*_*rs*_(λ) data collection, as well as SDP model application, are found in (30) and SDP model code is available on GitHub (see Data Availability Statement). Briefly, the concentrations of thirteen phytoplankton pigments are derived from the residual between a measured *R*_*rs*_(λ) spectrum and a modeled *R*_*rs*_(λ) spectrum. For both the modeled and measured pigments, the sum of the accessory pigments used here was highly correlated with total chlorophyll-*a* concentration (measured R^2^ = 0.98; modeled R^2^ = 0.97). Thus, the fraction of each accessory pigment normalized to the sum of those accessory pigments was used to compare composition in the surface ocean across samples in both basins.

## Acknowledgments

SJK was supported by a Simons Foundation Postdoctoral Fellowship in Marine Microbial Ecology (Award #986836). NASA grants 80NSSC21K0015 to MLE & CAD, 80NSSC18K1431 to AES, and 80NSSC17K0716 to TAR supported this work. SJK, CAD, and SS were additionally supported by the David and Lucile Packard Foundation. This work could not have been accomplished without the Captains and crews of R/V *Roger Revelle* and RRS *James Cook*, as well as support from all other members of the EXPORTS science team and particularly the Trap Team. We are especially grateful to Pat Kelly, Sean O’Neill, Melissa Omand, and Ken Buesseler.

## Data and Code Availability

All DNA sequence data, POC flux data, HPLC pigment data, and hyperspectral remote sensing reflectance data shown here can be found on the EXPORTS page of NASA’s SeaWiFS Bio-optical Archive and Storage System (SeaBASS): https://seabass.gsfc.nasa.gov/experiment/EXPORTS. Bulk sediment trap and individual particle DNA data can be found at: https://seabass.gsfc.nasa.gov/archive/MBARI/durkin/EXPORTS. Whole seawater DNA data can be found at: https://seabass.gsfc.nasa.gov/archive/URI/rynearson/EXPORTS/. Bulk POC flux data can be found at: https://seabass.gsfc.nasa.gov/archive/MAINE/estapa/EXPORTS/. Code for the SDP model can be found at: https://github.com/sashajane19/Rrs_pigments. Code for DNA data analysis can be found at: https://github.com/sashajane19/EXPORTS_particle_DNA. If you are unable to find any data products or code, please contact the corresponding author.

