## Supporting Information for "Ocean carbon export can be predicted from ocean color-based phytoplankton communities"

**Supplementary Methods:**

***Sediment trap deployment and sample treatment:***

Each trap platform had four collection tubes at each depth (diameter = 12.7 cm). Two of the tubes contained 0.3% formaldehyde-poisoned brine (70 ppt salinity) buffered to pH 8 with borate and overlain with 1 μm filtered surface seawater (1). Samples from these tubes were used for measurements of POC and other bulk quantities. POC analysis is described in detail in (1). Total particulate carbon was measured by combustion elemental analysis. Particulate inorganic carbon (PIC) was measured separately via coulometric titration (2) and POC fluxes were calculated by subtracting PIC from the total particulate carbon concentration. Uncertainty estimates were calculated from sample replicates.

The third tube contained either RNAlater (3) or 0.3% formalin and was used for bulk extraction of nucleic acids (see Supporting Information). A comparison of trap preservation methods (RNAlater vs. formalin) found that these samples had comparable Shannon diversity and eukaryotic community composition across methods, but bacterial community composition differed (Paul et al., *in prep*). The fourth tube contained a polyacrylamide gel overlain by 1μm filtered surface seawater (4, 5) and was used for measurements of nucleic acids in individual particles (6) and for quantitative particle classification and imaging (7). Upon trap recovery, particles collected in bulk collection tubes were allowed to settle for at least one hour before removing overlying seawater and draining the bottom sample layer. Samples were pre-filtered using a 330 μm mesh to separate zooplankton that swam actively into the sample. Zooplankton swimmers were visualized with a dissecting microscope and removed from the mesh (details in

(4)), and the remaining particles were rinsed back into the sample using filtered seawater.

Measured POC fluxes in the North Atlantic surface tethered traps (STTs) in the third deployment may have been subject to hydrodynamic effects, with evidence for under-collecting in the upper traps ((8); Estapa et al., *in prep*). However, both the bulk POC and DNA samples would have been affected by these hydrodynamic effects, so these paired samples are still used here for comparison.

#### *18S rRNA gene amplification and sequence analysis*

The V4 hypervariable region of the 18S rRNA gene was amplified using universal eukaryotic primers Reuk454FWD1 and V4r (9) on all sample types. Because seawater samples and particle samples were sequenced at different facilities, different Illumina-specific adaptors were added to the 5'ends of the primer used to amplify seawater DNA (forward5'- tcgtcggcatcagatgtgtataagagacagccagcascygcggtaattcc-3', reverse 5'-GTCTCGTGGGCTCGGAGATGTGTATAAGAGACAGACTTTCGTTCTTGAT-3') compared to the primers used to amplify particle DNA [ref 42] (forward 5'-acactgacgacatggttctacaccagcascycgcctaattcc -3' and reverse 5'-TACGGTAGCAGAGACTTGGTCTACTTTCGTTCTTGAT-3'). Seawater DNA was amplified in PCR reactions containing 1X AccuStart II PCR mix (VWR 89235-018), 0.3 µM each forward and reverse primer, and 20-100 ng DNA template using the following protocol: 2 min at 94°C, followed by 30 cycles of 94°C for 30 s, 55°C for 30 s, and 72°C for 1 min, and a final 10 min incubation at 72°C. Particle DNA was amplified in 25 µl PCR reactions using the KAPA Hifi Hotstart PCR kit (Roche KK2502) containing 1x buffer, 0.2 U polymerase, 0.3 mM dNTP, 0.3 mM each forward and reverse primer and 5 ml DNA extract with the following protocol: 5 min at 95°C, followed by 35 cycles 98°C for 30 sec, 62°C for 45 sec, and 1 min at 72°C.

PCR amplicons were cleaned with Ampure XP beads (Beckman Coulter, Brea, CA, USA) and quantified with the Qubit High Sensitivity DNA Assay Kit (ThermoFisher Scientific, Waltham, MA, USA). Amplicons from surface seawater samples were amplified for an additional five cycles to add Nextera indices and adaptors (Illumina, San Diego, CA, USA) and cleaned again with Ampure XP beads. These PCR products were pooled and quantified with the KAPA qPCR kit (Kapa Biosystems, Wilmington, MA, USA) prior to Illumina MiSeq sequencing with V3 chemistry (2 × 300 bp reads) at the University of Rhode Island Genomics and Sequencing Center. For sediment trap samples, barcoding and additional cleanup steps were performed at the Michigan State University Sequencing center prior to Illumina MiSeq sequencing with V3 chemistry (2 x 250 bp reads).

### Supporting Figures

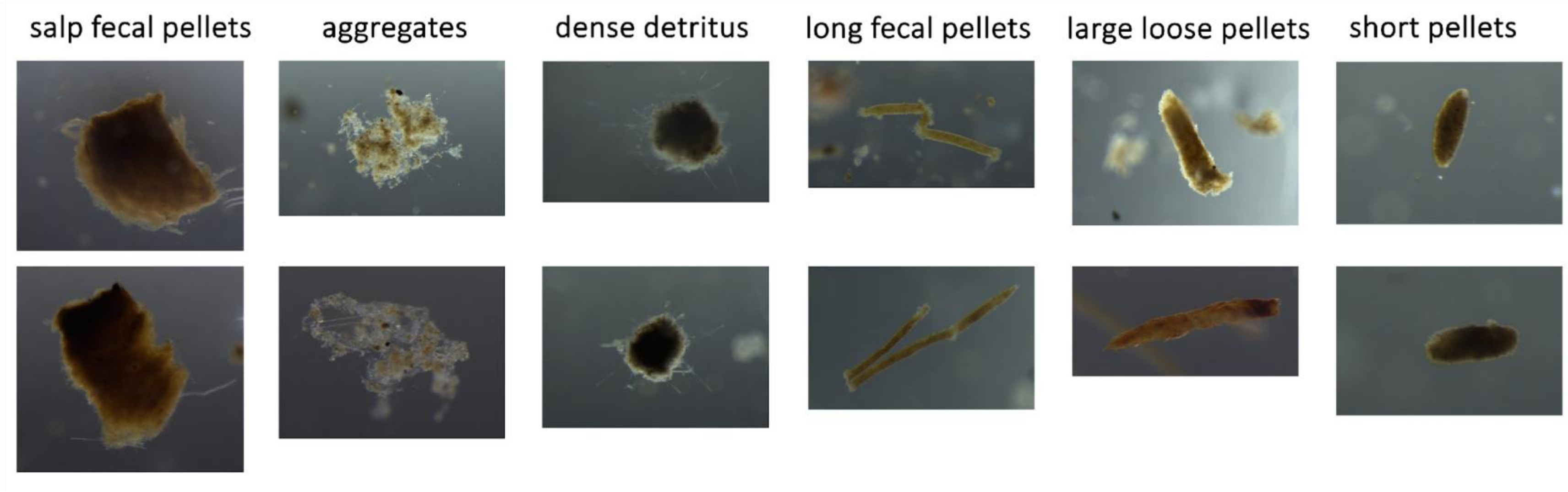


**Figure S1.** Example images for the major particle types described in this analysis.

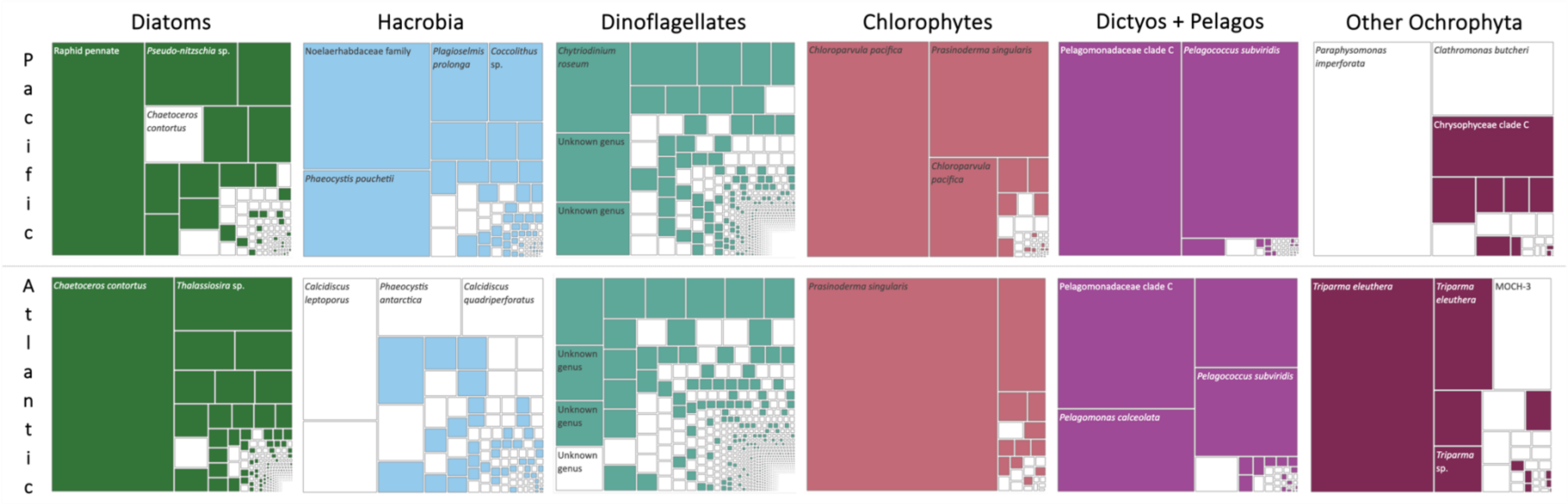


**Figure S2.** Major pigment-based phytoplankton group abundance in sinking particles (bulk particles + individual particles) in the North Pacific (top row) and North Atlantic (bottom row). Each small box within a larger box represents an individual ASV within that group. If the box is colored in, that ASV was found in a surface seawater sample. Boxes that are not colored in represent ASVs that were not found in the surface.

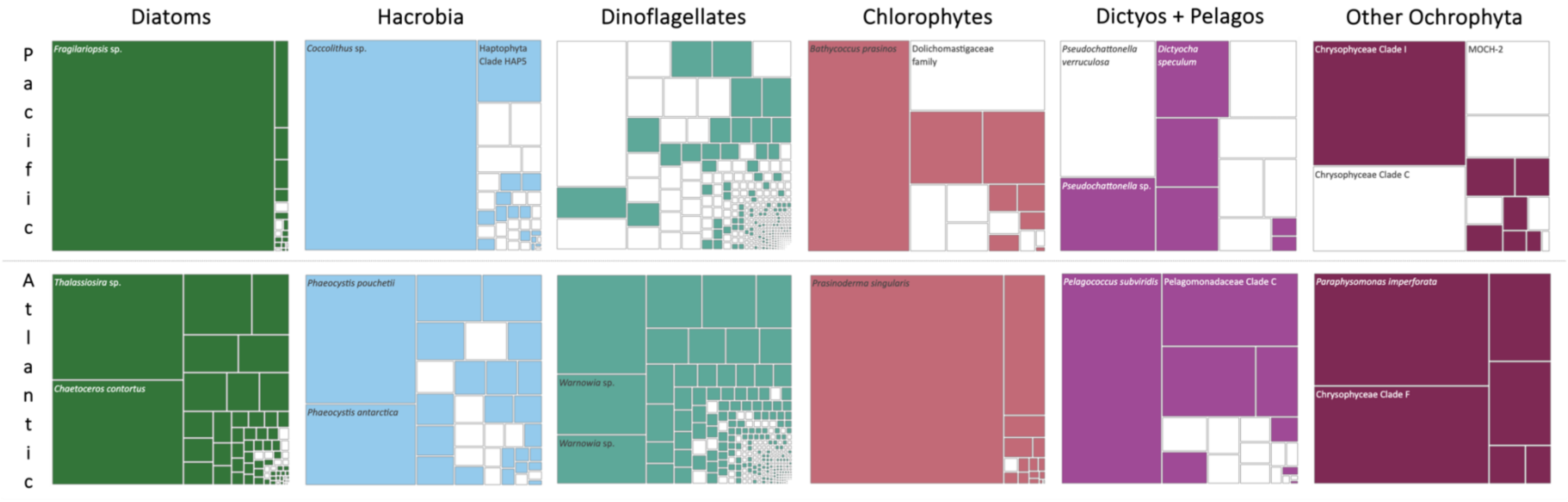


**Figure S3.** Major pigment-based phytoplankton group abundance in bulk sinking particles in the North Pacific (top row) and North Atlantic (bottom row). Each small box within a larger box represents an individual ASV within that group. If the box is colored in, that ASV was also found in an individual particle sample. Boxes that are not colored in represent ASVs that were found in bulk sediment trap samples but not in individual particle samples.

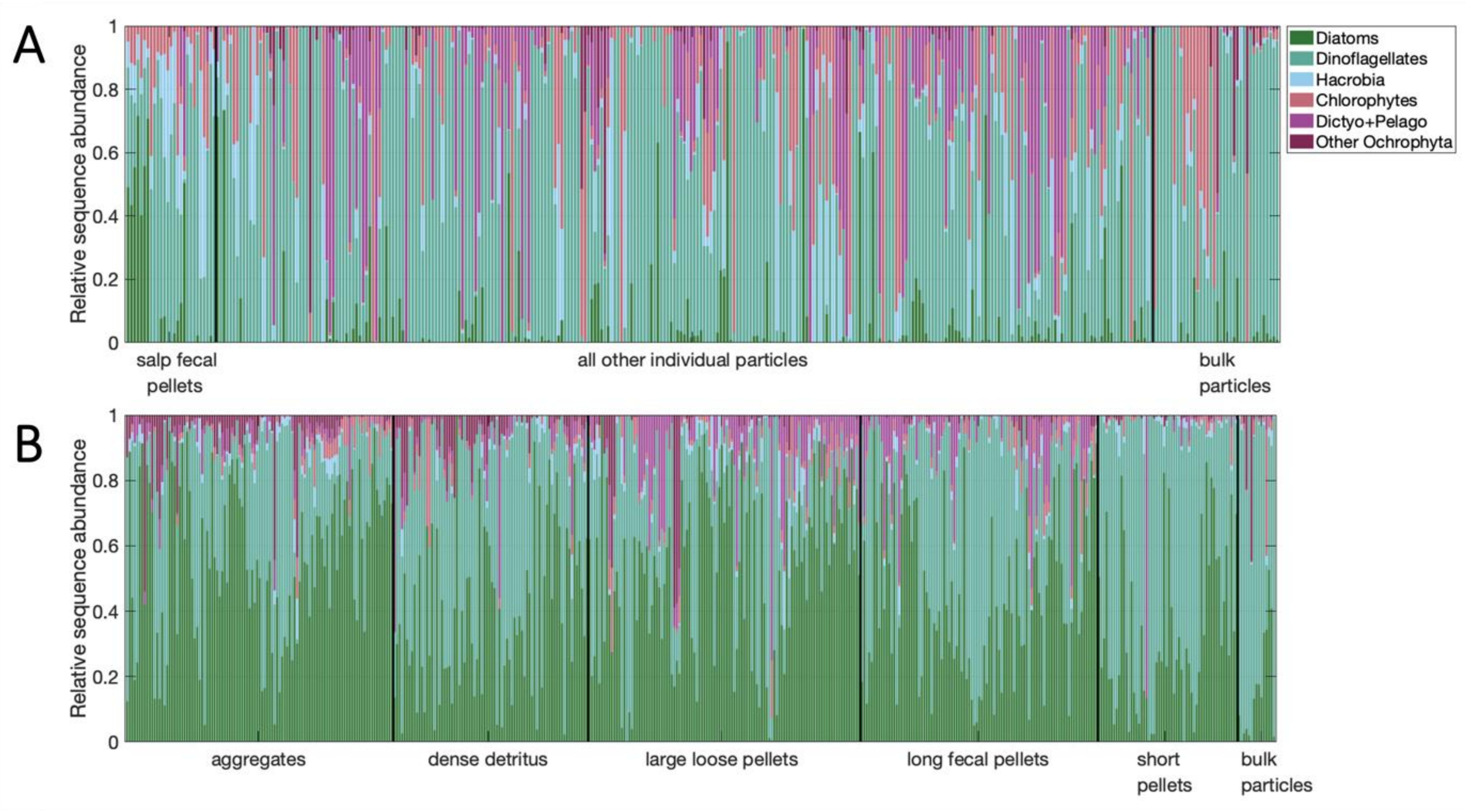


**Figure S4.** Particle composition across the six major phytoplankton groups shown here for all statistically different types of individual particle samples and all bulk particles samples in (A) the North Pacific and (B) the North Atlantic.

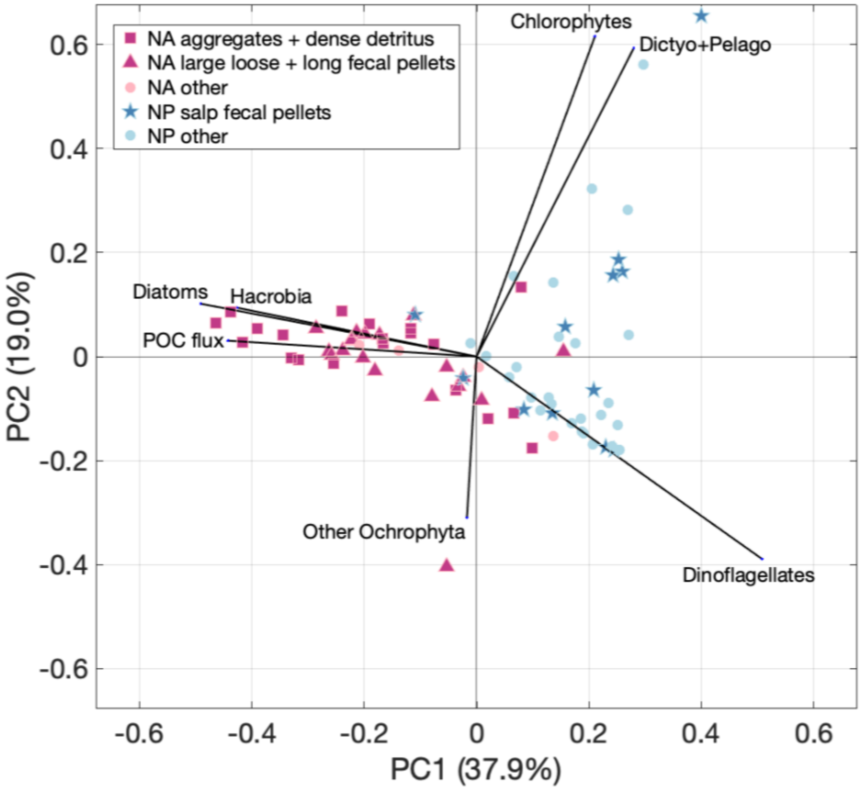


**Figure S5.** Principal components analysis (PCA) of particle-specific data. The orientation of each group of particles from each trap and each PCA input variable (black lines) is shown for the first 2 components (Component 1 = 37.9% of variability; Component 2 = 19% of variability).
